# Effects of order on memory of event times

**DOI:** 10.1101/2020.09.11.289058

**Authors:** Michelangelo Naim, Mikhail Katkov, Misha Tsodyks

## Abstract

Memorizing time of an event may employ two processes (i) encoding of the absolute time of events within an episode, (ii) encoding of its relative order. Here we study interaction between these two processes. We performed experiments in which one or several items were presented, after which participants were asked to report the time of occurrence of items. When a single item was presented, the distribution of reported times was quite wide. When two or three items were presented, the relative order among them strongly affected the reported time of each of them. Bayesian theory that takes into account the memory for the events order is compatible with the experimental data, in particular in terms of the effect of order on absolute time reports. Our results suggest that people do not deduce order from memorized time, instead people’s memory for absolute time of events relies critically on memorized order of the events.

## Introduction

Tulving [1, 2, 3] proposed a distinction between semantic memory (general knowledge) and episodic memory (personal experiences that carry information about time and location). We know very little about how time is encoded in the brain, hence theoretical understanding of episodic memory is difficult. Our introspective experiences and psychological studies indicate that event time can be remembered either in the absolute form (when the event happened) or in the relative form (whether the event happened before or after other specific events) [4]. Absolute time processing is quite reliable for short intervals, such as when catching a ball or playing an instrument, but deteriorates when longer intervals are involved, to the extent that we are often unaware of when some events happened (for example, one may know that Robert Kennedy was assassinated later than his brother was, but may not know when this assassination happened). Most of psychological studies of time memory relate to event duration rather than their absolute occurrence time (see e.g. [5, 6]).

In the current contribution, we consider the issue of interactions between absolute and relative time representations in episodic memory. In particular, lower-level absolute event time is a continuous feature while higher-level relative time order between events is of a discrete nature, hence one could expect that as time goes by, the latter could be more reliably stored in memory and hence take precedence in inference process. Interactions between absolute and relative attributes was indeed observed in recent studies of memory for simple visual attributes [7, 8]. In particular, reports of relative orientation of stimuli to reference strongly biased subsequent reports of absolute orientation of the stimulus [7]. In [8], multiple stimuli were presented to observers who did not have to explicitly report their relative orientations, only the absolute orientation of each stimulus. Still, these reports were strongly biased by the relative orientations between the stimuli. Moreover, the orientation reports could be predicted quantitatively by retrospective Bayesian inference of absolute orientations if relative orientations were assumed to be treated by the brain as given, i.e. decoding followed reverse hierarchical scheme from complex to simple features, as opposed to direct hierarchy of encoding (see also [9]). These intriguing results raise some fundamental issues on the nature of information encoding and decoding in the brain at the time when stimuli that give rise to perception are withdrawn.

Here we study how generic the retrospective decoding in memory is and whether it can be extended to the domain of event times in episodic memory. To this end, we performed experiments where either single events or sequences of several events were presented to participants at different times, after which they had to report their time of appearance. Following [8], we evaluated the interaction between absolute presentation time of each event and relative order between them. We observed very strong interference between these two types of information, such that reports of absolute time of an event were consistently shifted towards earlier or later times depending on the inferred order of that event relative to the other ones. We also developed Bayesian inference scheme for absolute time reports and compared it to our experimental observations.

## Experimental design and results

Experimental design is illustrated in the Fig. 1. Initially, we wanted to establish the quality of absolute time encoding of an event, when no other events were present during the same episode. Therefore, in the first experiment participants were exposed to a list of three items (words or images; see Methods for more details). Each trial was divided into 11 time slots of duration 1.5 seconds each. The first and the last item were always presented in the first and last slot, respectively, to delineate the beginning and the end of the trial, while the second item was presented in a randomly chosen intermediate slot. Each item was shown for 1000 ms in the beginning of a slot. The experiment was performed using Amazon Mechanical Turk ®. Participants were then requested to report the presentation time of one specific item, by moving a green circle with the mouse to the correct position on a sliding bar. At the beginning of each experiment, 5 training trials were performed where participants received a feedback with the correct timing (location of a circle) presented to them on another bar. Additional 15 trials without feedback were subsequently performed for data analysis. Results obtained in the first experiment are presented in Fig. 2 where reported time distributions for each presentation time of the intermediate item are shown. One can see that reported time distributions are rather wide, except for the very beginning, end and middle of a trial. Moreover, it is interesting that the results are very similar for both words and images.

**Figure 1:**
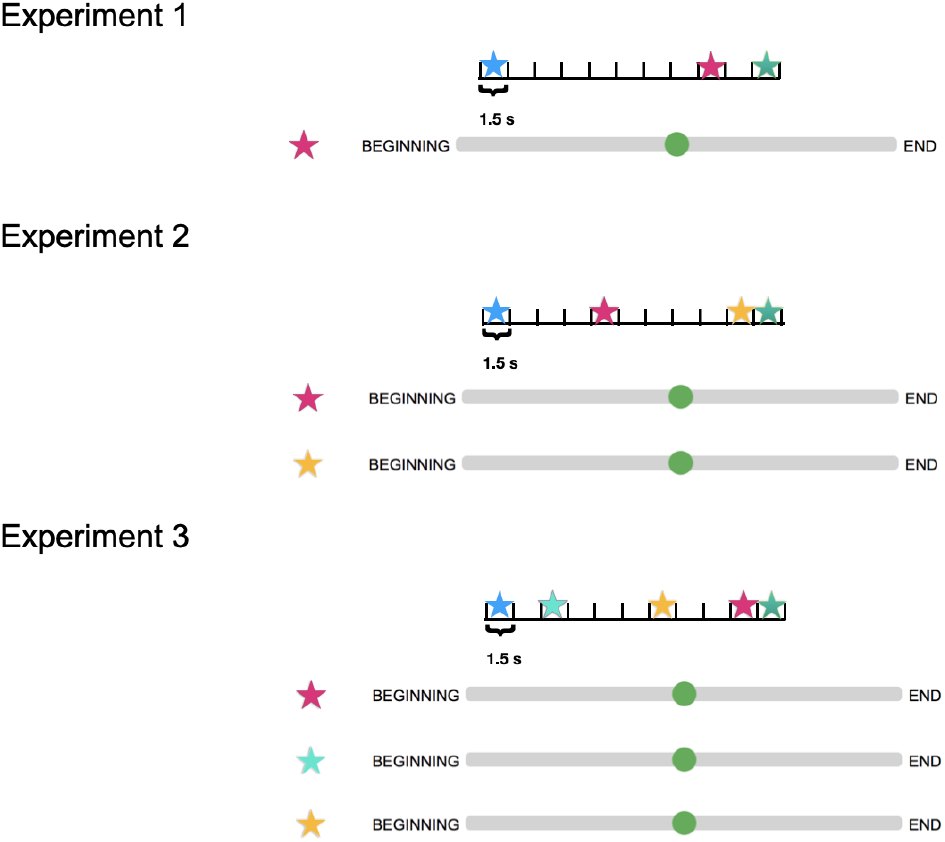
Experimental design. Upper panel: Experiment 1 scheme, three items presented. One item always presented at the beginning of the trial, one always at the end. The duration of a trial was divided into 11 slots of equal size. Intermediate stimulus was presented in a randomly chosen slot. After the presentation participants have to report the time of one of the items by moving a green cursor to the corresponding position on a sliding bar. Middle panel: Experiment 2 scheme, four items presented. One item is always presented at the beginning of the trial, another one at the end of the trial, while two intermediate items are presented in random time slots. After presentation participants have to report the time of two intermediate items by moving green cursors to the corresponding positions on two sliding bars. Vertical location of earlier and later items were random. Lower panel: Experiment 3 scheme, five items presented. One item is always presented at the beginning of the trial, another one at the end of the trial, while three intermediate items are presented in random time slots. After presentation participants have to report the time of three intermediate items by moving green cursors to the corresponding positions on three sliding bars. The correspondence between vertical location of a sliding bar and item was random.

**Figure 2:**
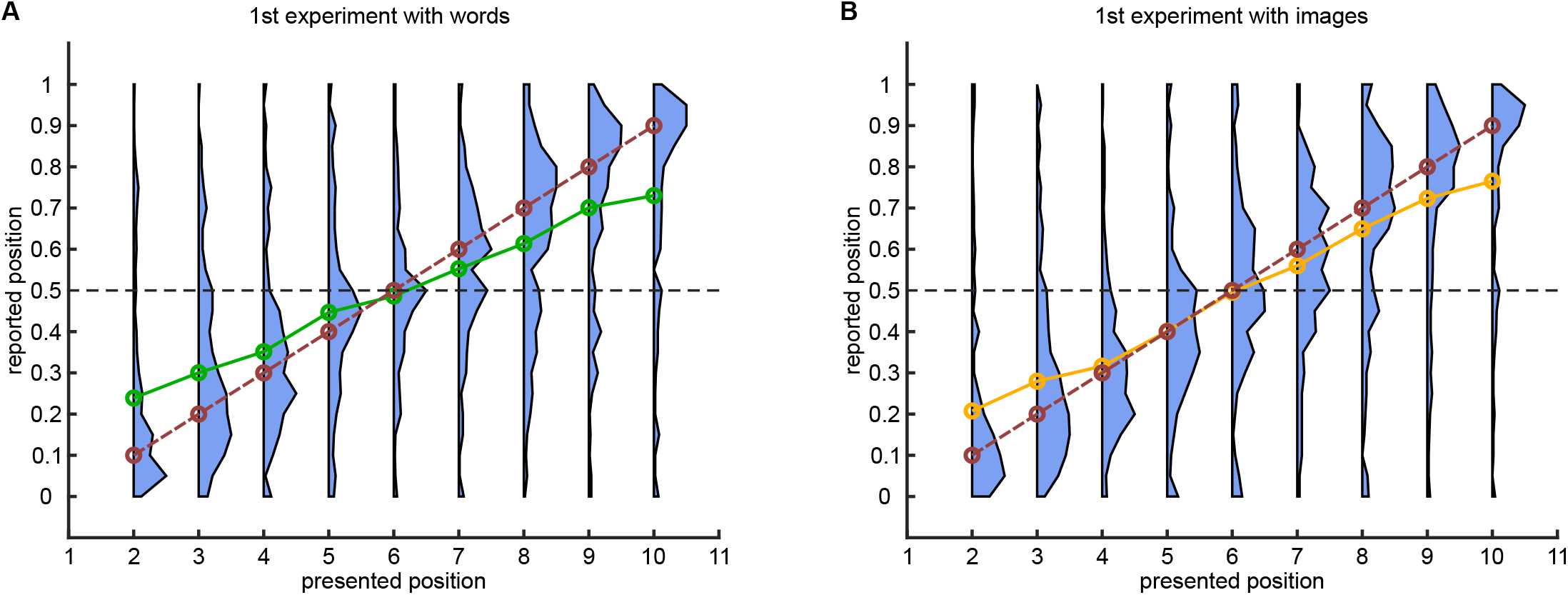
Experiment 1: distribution of reported times. **(A)**: For each presentation time of a word distribution of reported times. Green line corresponds to average of the distributions, dashed red line corresponds to perfect report. **(B)**: Same for images, where the orange line corresponds to average of the distributions.

To test the effect of event order on memory of “absolute” time, the second experiment with lists of four items was performed. Participants were requested to report the time of two of them, see Fig. 1.

Before analyzing Experiment 2, it is instructive to form a prediction for the accuracy of relative time order for two intermediate items based on the results of Experiment 1. If we assume that two intermediate items are encoded and reported independently, we can predict from Fig. 2 the probability that for any two presentation times the participants will make a mistake in ordering the items (see Figs. 3A and 3D). As expected, when the two presented items are close to each other, the predicted probability to flip the order is higher. However, experimental results do not show this tendency, with flip probability being rather uniform across all presentation conditions (see Figs. 3B and 3E). Overall, the accuracy of time ordering was 93% for words and 92% for images as opposed to 75% and 77% as predicted from the results of Experiment 1. The measured values are significantly different from predicted ones with *p* ≈ 10^−90^ and *p* ≈ 10^−54^ for words and images respectively (Fisher exact test). These results show that order of events in memory is not derived from independent estimates of timing for each of the events, suggesting that it is stored separately from them.

**Figure 3:**
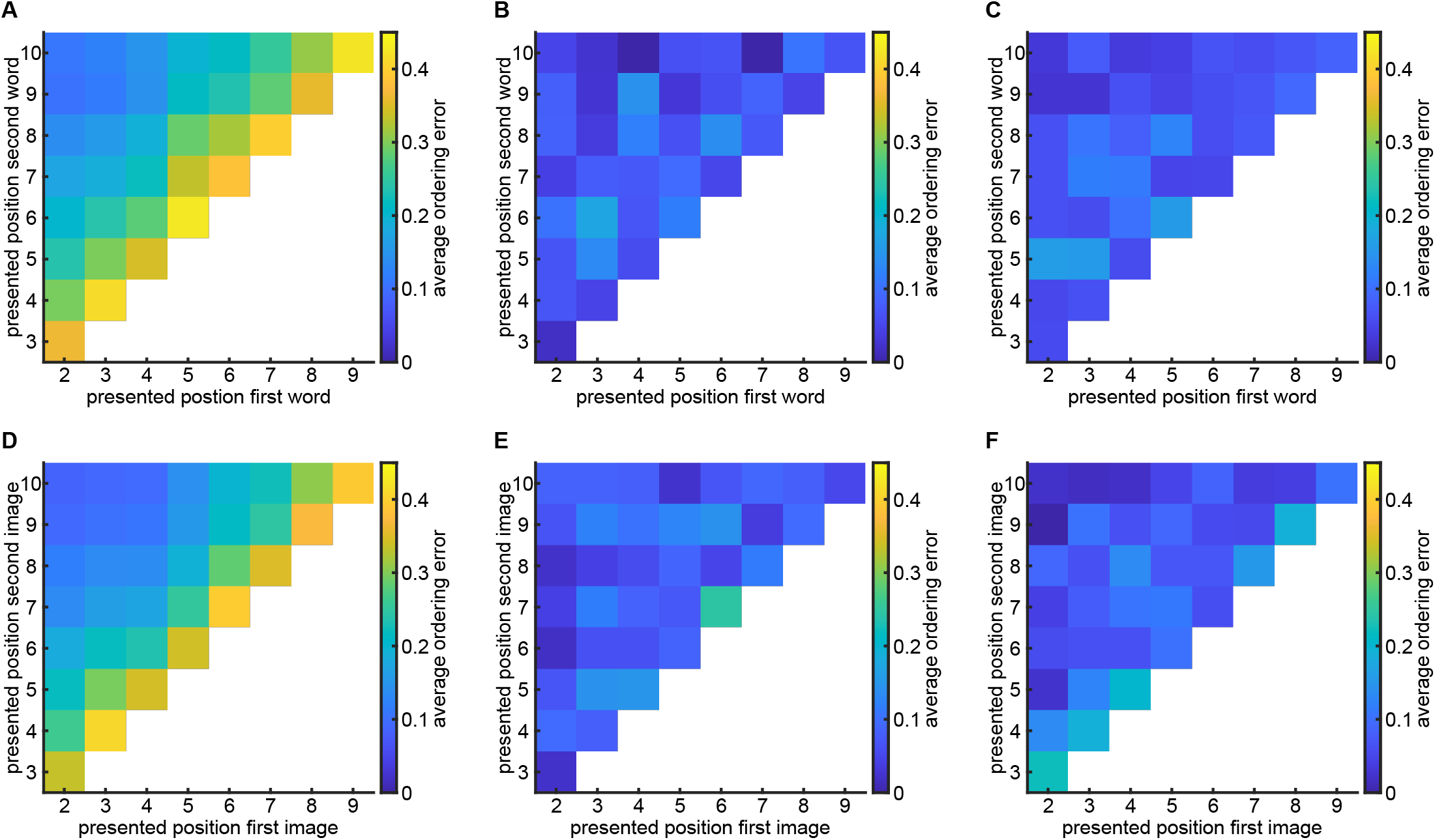
Accuracy of relative time ordering in Experiment 2. **(A)**: Naïve prediction of average ordering error from independent distributions obtained in the Experiment 1 with words. **(B)**: Experimental average ordering error with two presented words. **(C)**: Bayesian simulations of average ordering error. **(D - F)**: Same as (A - C) for images.

To probe the interactions between ordinal and absolute time representations in memory, we analyzed absolute time reports for the first and second items separately. Fig. 4 shows the average reported time as a function of presented time for both first and second items (red an blue dashed lines, respectively). One can see a clear and significant effect of order on absolute time reports, with participants tending to report the first intermediate item earlier than the second item on the trials when they are presented at the same time slot. The difference between the average report times of the first and second items, for the same presentation time, is significant for all time slots, except last time slot for images where much fewer trials were collected. Indeed, comparing reported times for the first or second stimulus presented in different trials on the same temporal bin using t-test with unequal variance lead to *p* < 0.01, *p* = 0.04 and *p* < 10^−4^ when words are presented on second, 9^th^ and all the rest time slots respectively; *p* = 0.06, *p* = 0.013 and *p* < 10^−4^ when images are presented on second, 9^th^ and all the rest time slots respectively. Still the average report time depends on the presented time for both cases, indicating a complex interaction between the ordinal and absolute information in memory, which will be considered in more details in the next section. The corresponding report histograms are shown in Fig. S1.

**Figure 4:**
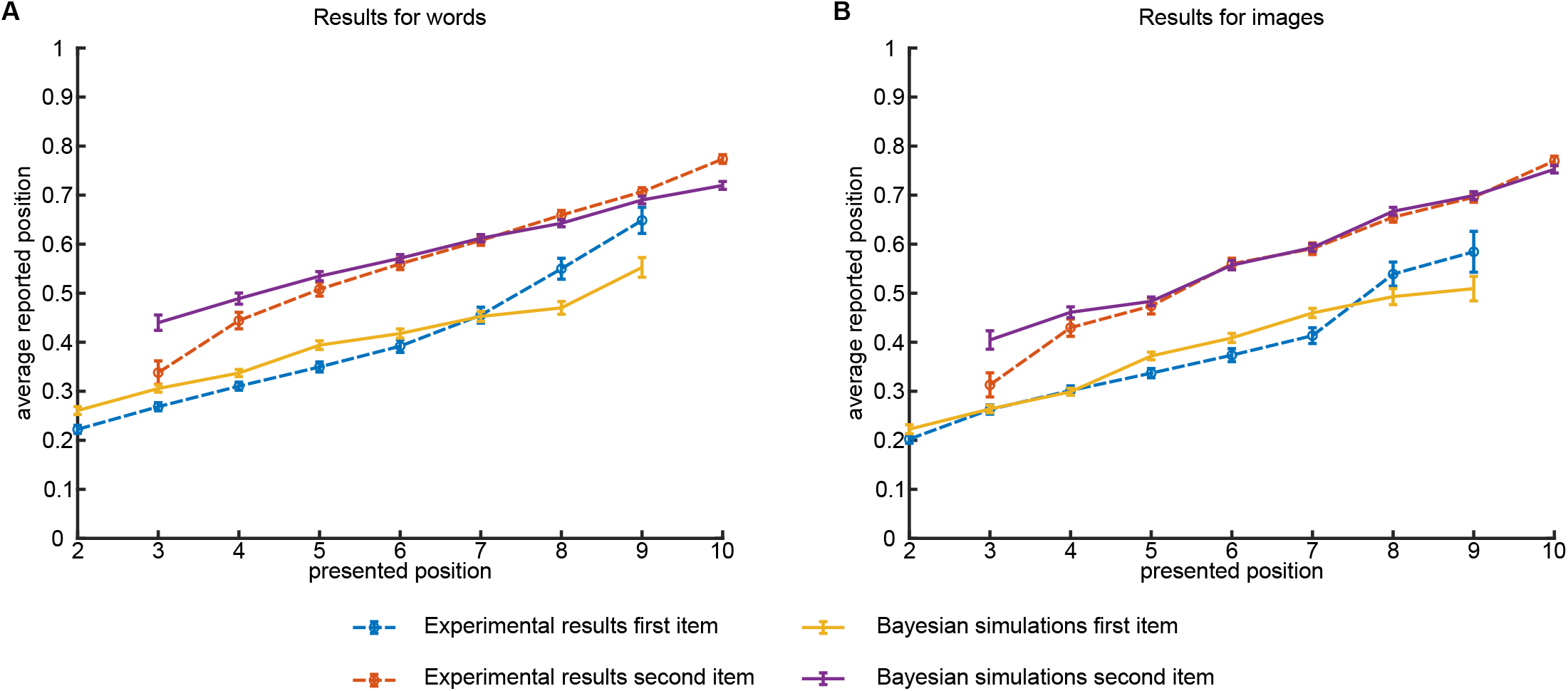
Experiment 2: average time reports. **(A)**: Average report times for first and second word, separately. Dashed lines: experimental results; solid lines: results of Bayesian inference **(B)**: Same as (A) for images.

To further evaluate the interactions between ordinal and absolute time representations in memory, we conducted the experiment with three intermediate items (experiment 3, see Fig. 1). In this case, the participants could either report the items in the correct order or make one of five possible ordering mistakes, see Fig. 5. Also in this case, the overall probability to report the correct ordering is still high: 77% (leftmost red column), which is significantly better than predicted from experiment 1 assuming independent reports (left-most blue column; 37%, *p* ≈ 10^−300^, exact Fisher test). This result is consistent with classical recall experiments showing that short lists of words can be well recalled in the correct order (see for example [10]). The effect of ordering on absolute time reports is even stronger for three items, for example a shift of reports for first and last intermediate items relative to their actual positions is bigger if there is another item presented between them. In other words, the remembered time interval between past events is larger when other events intervene between them, which is consistent with previous work [11]. However the reported times still depend on the presented ones, albeit with a smaller slope than in the case of two items (Fig. 6, blue, red and green dashed lines for first, second and third item, respectively). In particular, if one considers a linear model for the reported position with presentation position as a factor and consider a null hypothesis that reported position does not depend on the presentation one, the null hypothesis can be rejected (*F* (2233) = 21.1, *p* = 5 · 10^−6^, *F* (4468) = 781, *p* = 2 · 10^−158^, *F* (6703) = 3780, *p* = 0, for first, second and third words).

**Figure 5:**
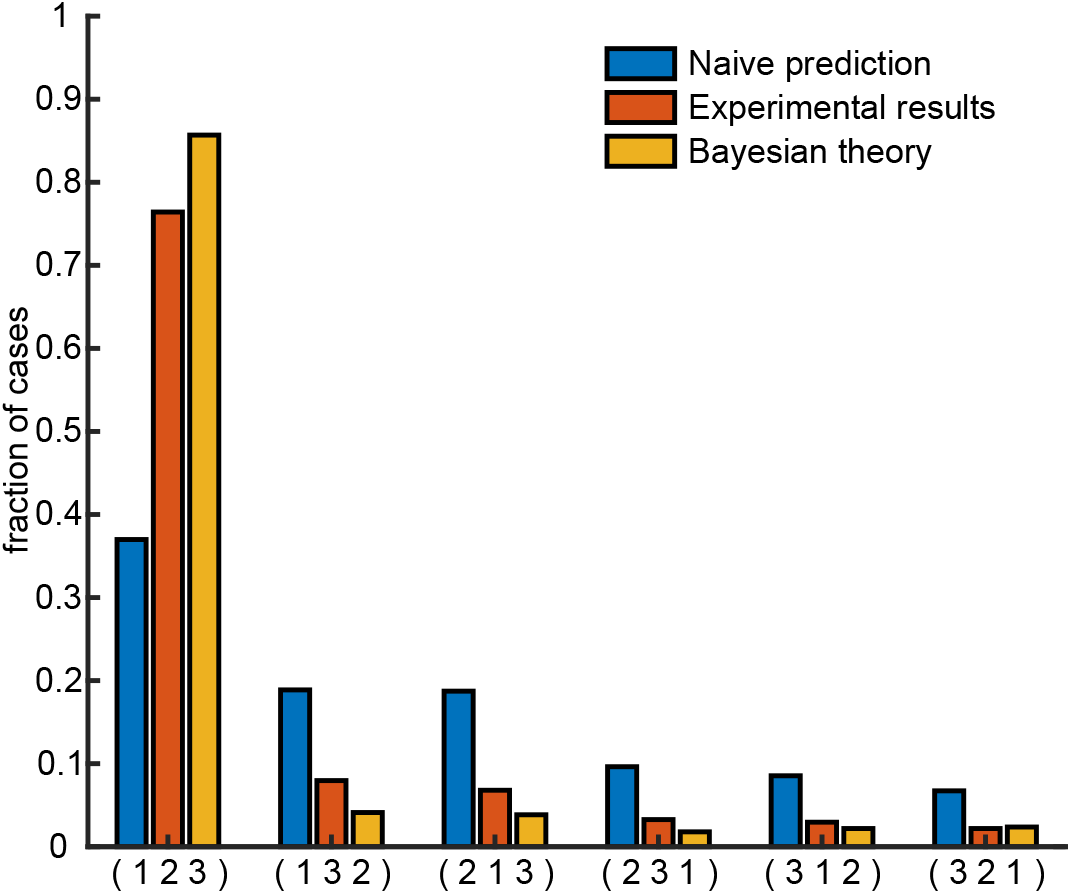
Experiment 3: Accuracy of time ordering. Probability of each of 6 possible ordering permutations of the 3 presented items, where 1,2,3 stand for first, second and third presented items, respectively. Blue columns: Naïve prediction from independent samplings of distributions obtained in the experiment 1 with words. Orange columns: experimental results. Yellow columns: Bayesian theory.

**Figure 6:**
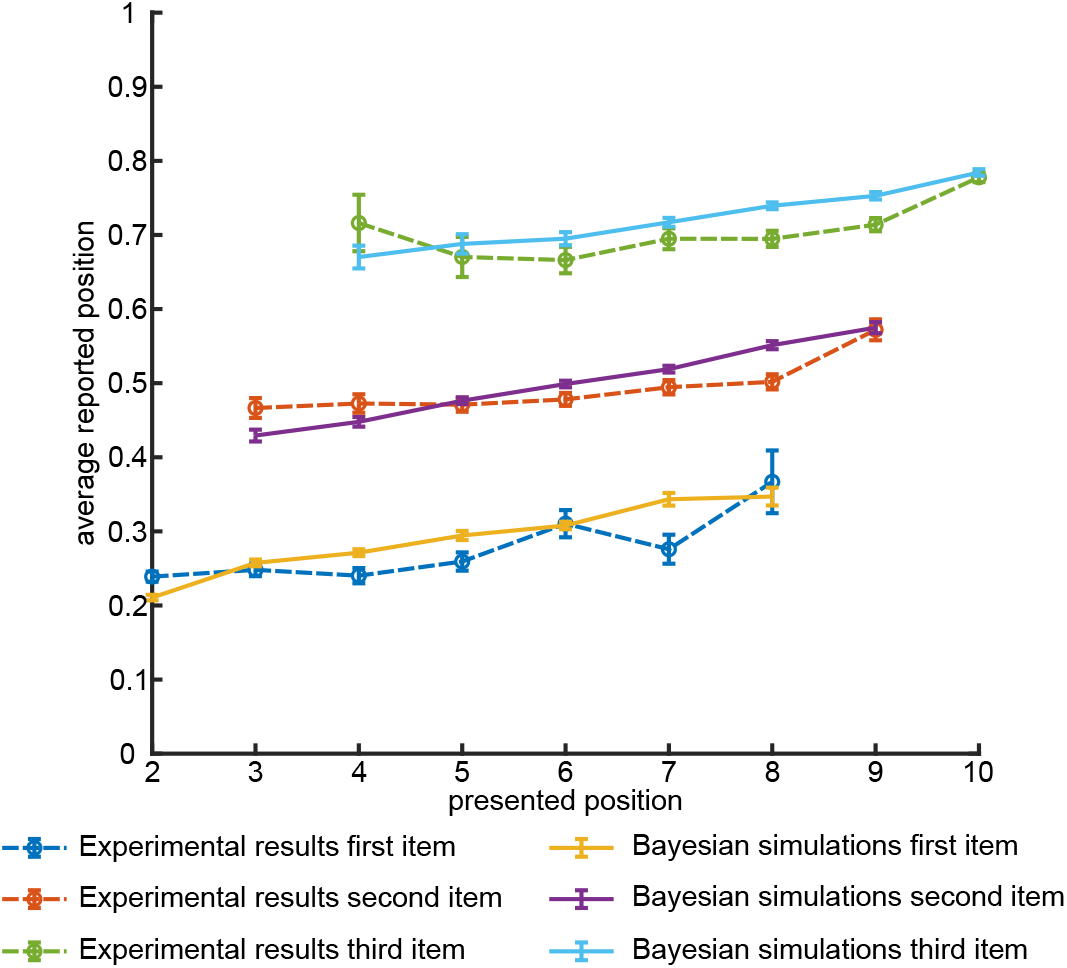
Experiment 3: average time reports. Average report times for first, second and third word, separately. Dashed lines: experimental results; solid lines: results of Bayesian inference

## Bayesian Theory

The results presented above indicate that absolute and relative times are two interactive but distinct aspects of episodic memory. We therefore developed a Bayesian time decoding theory that elaborates the precise nature of this interaction. Here we present the theory for the case of two items (see Methods for more details). Define *t*_1_ and *t*_2_ the absolute presentation times of events within a trial, and 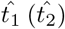 their internal representations at the report time. Define also *σ* = 1(0) if *t*_1_ < *t*_2_ (*t*_1_ > *t*_2_), respectively, to be the relative order between the events, and 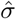 its internal representation. A crucial assumption of our theory is that continuous representations 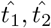 and discrete representation 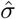 are stored independently in memory and have their own representational errors, hence 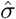 is not necessarily consistent with 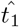 and 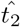.

The likelihood function for the internal variables is given by

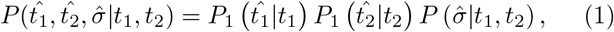

where we assume that 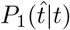 can be evaluated from the report time distribution measured in Experiment 1 (see Fig. 2) and

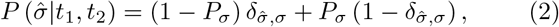

where *δ_k,m_* is a Kronecker delta

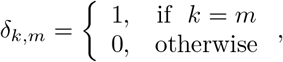

and *P_σ_* is the probability of a mistake in the internal representation of a relative order between events.

We assume that once all relevant features (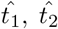 and 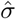) are represented in working memory, at the report time the brain performs Bayesian inference of absolute presentation times according to

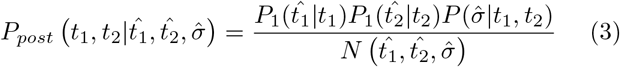

where 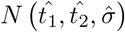 is the normalization and we assumed that prior distribution of event times is uniform in agreement with the experimental protocol (see Methods). One can see from Eqs. (2) and (3) that as the estimation of presentation order becomes more precise (*P_σ_* → 0), it serves as an effective prior for the estimates of the absolute presentation times. Following [8], we assume that reported times for the first and second items (*t*_1*r*_ and *t*_2*r*_, respectively) are generated as averages over the posterior distribution:

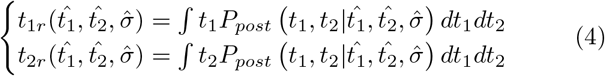

Since 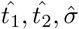 are distributed according to the likelihood function of Eqs. (1) and (2), this equation gives rise to the distributions of reported times *t*_1*r*_, *t*_2*r*_ for given presented times *t*_1_, *t*_2_. To generate these distributions, we randomly sampled the reported times in Experiment 1 as a proxy for 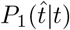, and used Eq. (2) to generate samples of 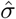. Using these samples, we formed the theoretical predictions for ordering mistakes probabilities (Figs. 3C and 3F) and average report time versus presentation time for both first and second item (Figs. 4A and 4B, overlaid with experimental results). The agreement between these predictions and the experimental results is rather good except for early and late positions (3rd position for the second stimulus and 9th position for the first stimulus). We speculate that when two intermediate stimuli are presented consecutively and immediately after the beginning of the trial (or before the end of it), participants form a separate memory for this sequence of three equally spaced stimuli and hence their reports are more reliable than what is predicted by the Bayesian model that does not include this mechanism. This effect disappears when we introduced a delay of 16 sec between the end of presentation and reports (see Experiment 4 in methods, and Figs. S6 and S7 for results). We also note that some of experimental report distributions exhibit bimodal shape that is not well captured by the model (Fig. S2).

The theoretical predictions for the Experiment 3 with three intermediate items were obtained as well and shown in Figs. 5 and 6 (see Methods). Also in this case Bayesian theory captures well the experimental results, most importantly the strong effect of ordinal information on absolute time representations in memory. The distribution of reported times and comparison with Bayesian model are presented in Figs. S3 and S4. A closer look at the experimental report distributions (Fig. S3) reveals that they contain a significant component that does not depend on absolute presentation time but only on the ordinal position of the corresponding item. This suggests that in this experiment, some people completely disregard absolute presentation times and only retain the temporal order of different items, i.e. their memory representation of time is quantified. Indeed, when we simulated our Bayesian model with zero ordering error *P_σ_* = 0 and uniform likelihood for absolute time 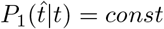 (see Eq. (1)), the obtained report distributions match well this component of the experimental distributions (see Fig. S5). We speculate that transition between gradual and quantized time representations happen when participants reach their working memory capacity for continuous variables like absolute time.

The precise nature of memory encoding of ordinal information is not clear at this stage. One possibility could be that rather than forming the explicit representation of presentation order, participants actively rehearse the presented items using working memory and use that at the time of absolute time reports. We therefore performed an additional series of experiments with one or two intermediate items where participants had to perform 16 seconds of math problems between the presentation of the items and time reports (Experiment 4). We reasoned that while performing calculations, participants would not be able to continuously rehearse the items. Nevertheless, we observed better-than-predicted precision of ordinal information in time reports in these experiments, as well as strong effects of order on absolute time reports (see Figs. S6 and S7), indicating genuine encoding of ordinal information in memory.

## Discussion

In this contribution we showed that absolute time of different events is not reliably represented in memory, while presentation order is. Moreover, the ordinal information strongly effects absolute time reports by shifting reported times according to the presentation order, even though ordinal information itself did not have to be explicitly reported by the participants in our experiments. Experimental results can be reasonably approximated by the Bayesian inference framework. These results are quite analogous to those of [8] in the visual domain. We therefore believe that they reflect a general principle in information processing according to which those aspects of information that are more reliably represented in memory take precedence to less reliably represented aspects and moreover act as Bayesian priors for inferring the latter. In particular, it appears that higher-level features such as ordinal relations between elementary components of complex stimuli that are of a discrete nature are decoded first and then constrain the decoding of lower-level, continuous features such as absolute time of an event or an absolute orientation of a line. The generality of the observed effects, in particular when longer time intervals are involved, should be investigated in future experiments.

## Acknowledgements

This research has received funding from the European Union’s Horizon 2020 Framework Programme for Research and Innovation under the Specific Grant Agreement No. 785907 (Human Brain Project SGA2); EU-M-GATE 765549 and Foundation Adelis. We thank Francesca Strappini for providing the images used in this study.

## Methods

### Participants, items and procedure

In total 1597 participants, were recruited to perform memory experiments on the online platform Amazon Mechanical Turk^®^ (https://www.mturk.com). Ethics approval was obtained by the IRB (Institutional Review Board) of the Weizmann Institute of Science. All experiments were performed in accordance with relevant guidelines and regulations. Each participant accepted an informed consent form before participation and was receiving 85 cents for approximately 10 min. For 1206 participants the presented lists were composed of non-repeating words randomly selected from a pool of 751 words produced by selecting English words [12] and then maintaining only the words with a frequency per million greater than 10, from Medler [13]. For 400 participants the presented lists were composed of non-repeating images (out of 149 possible): visual stimuli consisted of animal pictures [14], houses and body parts (free-copyrights Google images). Examples of the images are shown in Fig. 7. All the images were resized in browser to have width of 600 pixels. The items were presented on the standard Amazon Mechanical Turk^®^ web page for Human Intelligent Task (HIT). Each trial was initiated by the participant by pressing “Start Experiment” button on computer screen. List presentation followed 300 ms of white frame. During a trial, depending on the task, 3 to 5 items where shown in a total time frame of 16.5 seconds. More specifically, the trial was divided into 11 slots of 1.5 seconds each, and an item was shown in one of the slots. The first item was always presented in the first slot, the last item was presented in the last slot. Intermediate items were shown in randomly chosen slots with uniform probability. Each item was shown within HIT frame with black font at onset of slot for 1000 ms followed by empty frame for 500 ms. After the last item, there was a 1000 ms delay before participant performed the task.

**Figure 7:**
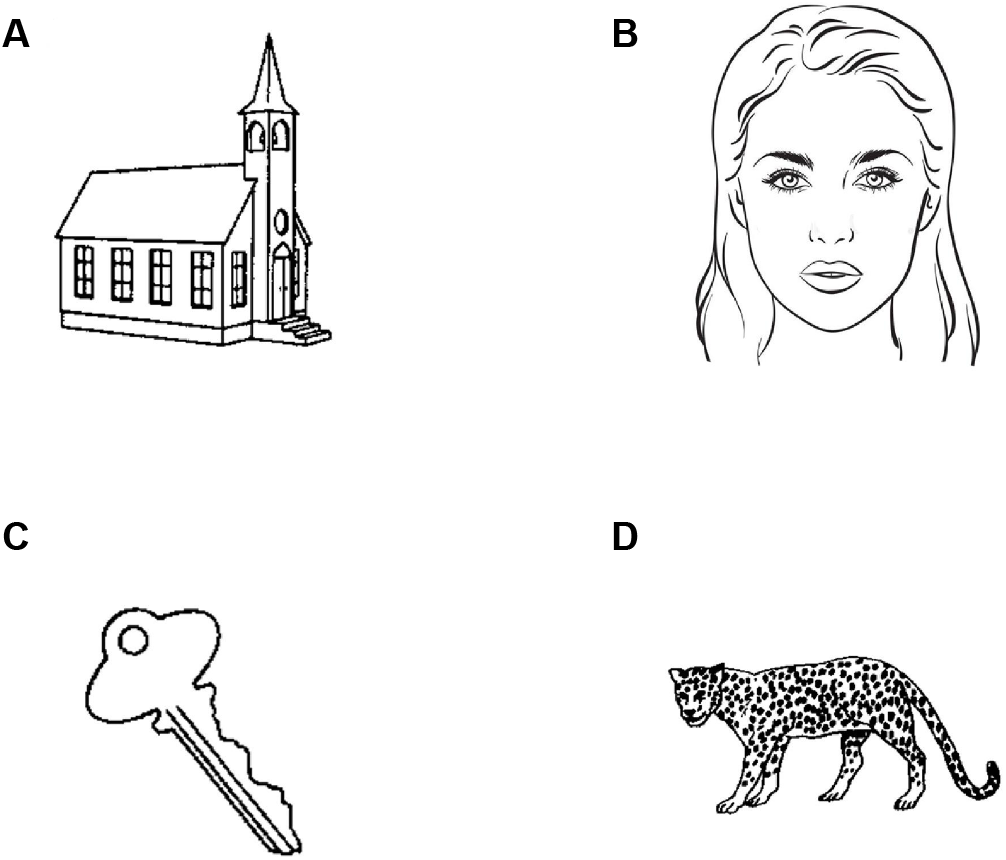
Images used in the experiments. **(A-D)**: four example images.

#### Experiment 1 - One intermediate item

Participants were presented with three items. One item was always presented at the beginning of the trial, one always at the end and one in a random slot. At the end, participants were requested to report the time of one of the items moving a green circle with the mouse to the correct position on a sliding bar. At the beginning there were training sessions where participants received a feedback with the correct timing (5 trials), followed by 15 trials without feedback. For 791 participants the items were words, for 400 the items were images.

#### Experiment 2 - Two intermediate items

Participants were presented with four items. One item was always presented at the beginning of the trial, one always at the end and the two others in random slots. At the end, participants were requested to report the time of two of the items moving a green circles with the mouse to the correct position on two sliding bars. Sliding bars corresponding to two items were presented vertically in randomized order. At the beginning there were training sessions where participants received a feedback with the correct timing (5 trials), followed by 15 trials without feedback. For 222 participants the items were words, for 198 the items were images.

#### Experiment 3 - Three intermediate items

451 participants were presented with five items. One item was always presented at the beginning of the trial, one always at the end and the three others in random slots. At the end, participants were requested to report the time of three of the items moving a green circles with the mouse to the correct position on three sliding bars. Sliding bars corresponding to three items were presented vertically in randomized order. At the beginning there were training sessions where participants received a feedback with the correct timing (5 trials), followed by 15 trials without feedback.

#### Experiment 4 - One and two intermediate items with longer delay

99 participants were presented with three items and 156 with four items. One item was always presented at the beginning of the trial, one always at the end and the others in random slots. At the end, participants had to perform 16 seconds of mathematical problem solving task and after that were requested to report the time of one or two of the items moving a green circles with the mouse to the correct position on the sliding bars. Sliding bars were presented vertically in randomized order. At the beginning there were training sessions where participants received a feedback with the correct timing (5 trials), followed by 15 trials without feedback.

### Participants selection

Examining the performance of each participant we encountered that many participants did not perform the task. For instance, since there are 6 possible orderings of items in Experiment 3, the chance to obtain correct order with random responses 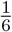, and since each participants performed 15 trials the expected number of correct trials from random responses is 2.5. In Experiment 2 there are only 2 orderings and the expected number of correct orderings from random responses is 7.5. As shown in Fig. 8, there is a bimodal distribution of the number of correct ordering in Experiment 3, with one mode around the chance performance. Therefore, we decided to exclude participants that had 6 or less correct orderings in Experiment 3 and less than 10 correct orderings in Experiment 2. After exclusion, we analyzed data for 190 participants in Experiment 2 with words, 181 with images, 256 in Experiment 3 and 173 in Experiment 4.

**Figure 8:**
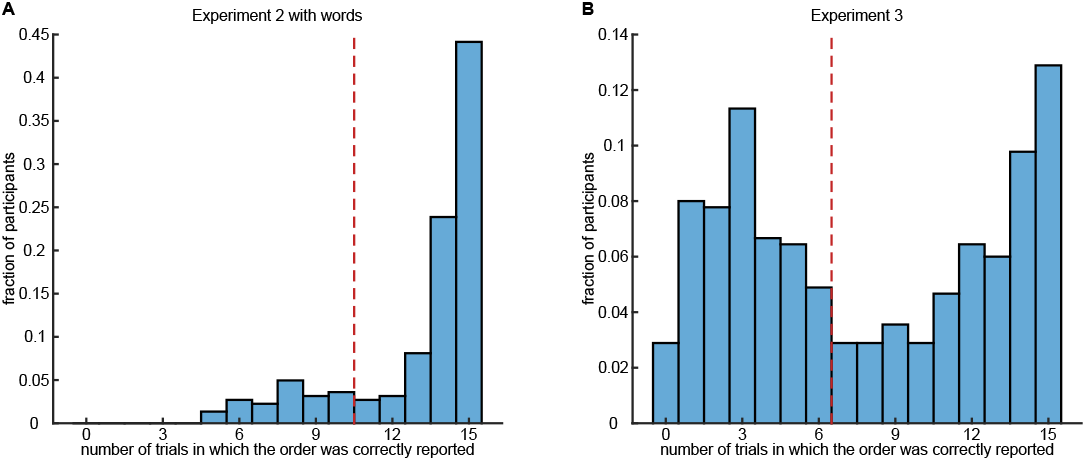
Participants selection. Distribution of fraction of trials in which the order was correctly reported for Experiment 2 with words **(A)** and Experiment 3 **(B)**. The dashed red line corresponds to the threshold used.

For Bayesian analysis:

- out of 791 participants that performed Experiment 1 with words, we used data only from the 190 participants that performed also Experiment 2.
- out of 400 participants that performed Experiment 1 with images, we used data only from the 181 participants that performed also Experiment 2.
- in Experiment 3, instead, a new pool of participants was used. Among them, only 34 previously completed Experiment 1. Therefore we considered all the 791 participants for the analysis.

Experimental data presented in this paper are available at https://osf.io/r3z9e [15].

## Bayesian inference with arbitrary number of presented items

To generalize the Bayesian equation for the case of *n* items in memory, we introduce a variable *σ* that denotes the ordering of their presentation times *t*_1_,…,*t_n_* (i.e. *σ* = (123) if *t*_1_ < *t*_2_ < *t*_3_ etc). We then write the likelihood function in the following form:

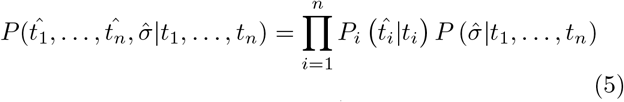

where we again assume that 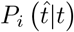 can be evaluated from the report time distribution measured in Experiment 1 (see Fig. 2). For different ordering likelihood functions, 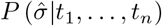, we assume for simplicity that

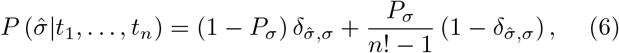

i.e. we assume that each ordering mistake in memory happens with equal probability. The rest of the calculations can then proceed as in *n* = 2 case presented in the paper. The value of *P_σ_* for simulating experiments 2 and 3 was chosen as 0.08 and 0.1, respectively. These values give rough correspondence between simulated and experimental reported positions and ordering errors.

## Author contributions

M.N. and M.K. analyzed the data, developed the mathematical model, wrote the manuscript text and edited the figures. M.T. has mentored and guided the project working on the data analysis, mathematical model and writing the manuscript text.

## Supplemental information

**Figure S1:**
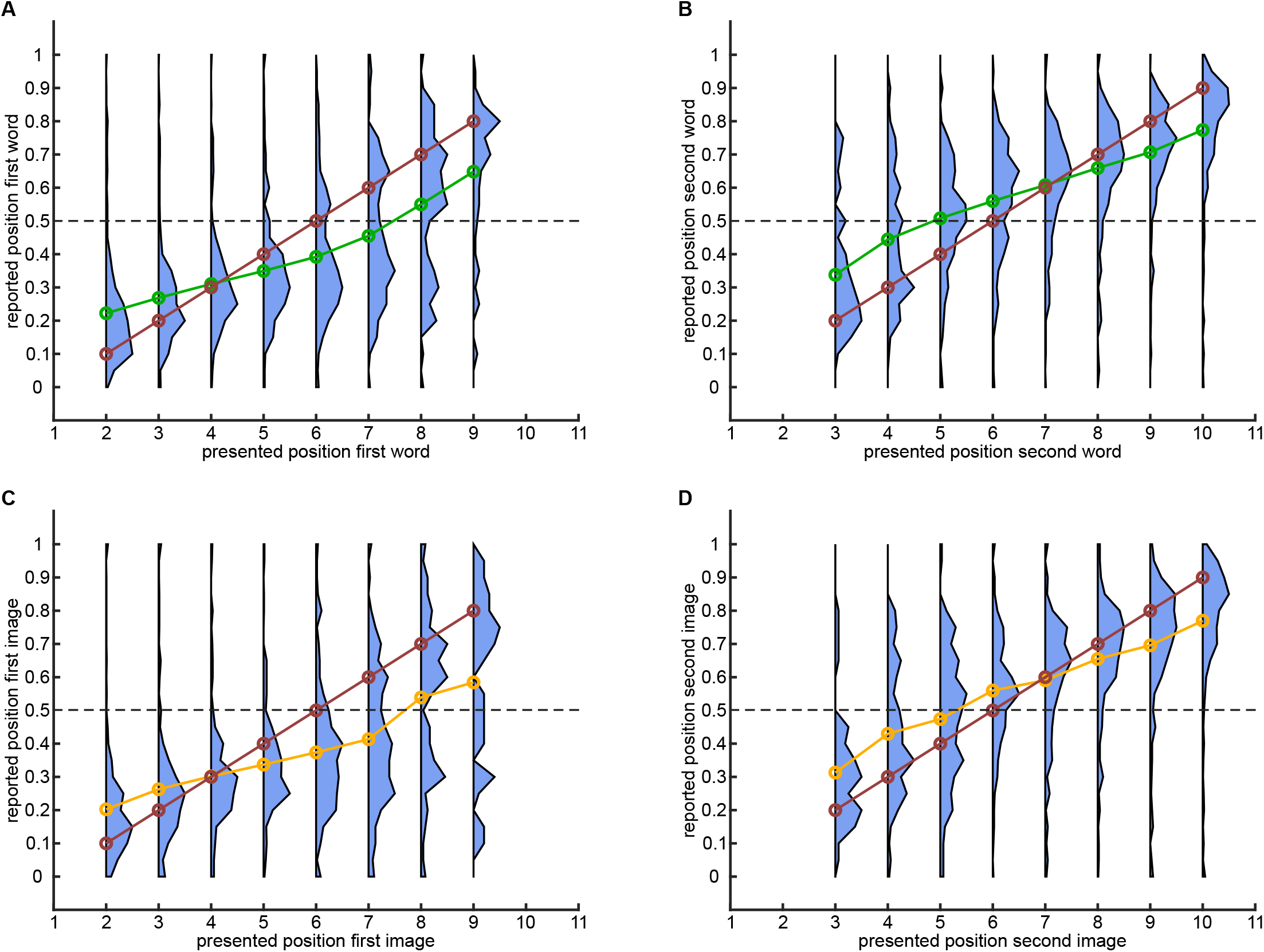
Experiment 2: distribution of reported times. **(A)**: For each presentation time of first intermediate word distribution of reported times. Green line corresponds to average of the distributions, dashed red line corresponds to perfect report. **(B)**: Same for second intermediate word. **(C)**: Same for first intermediate image, where the orange line corresponds to average of the distributions. **(D)**: Same for second intermediate image.

**Figure S2:**
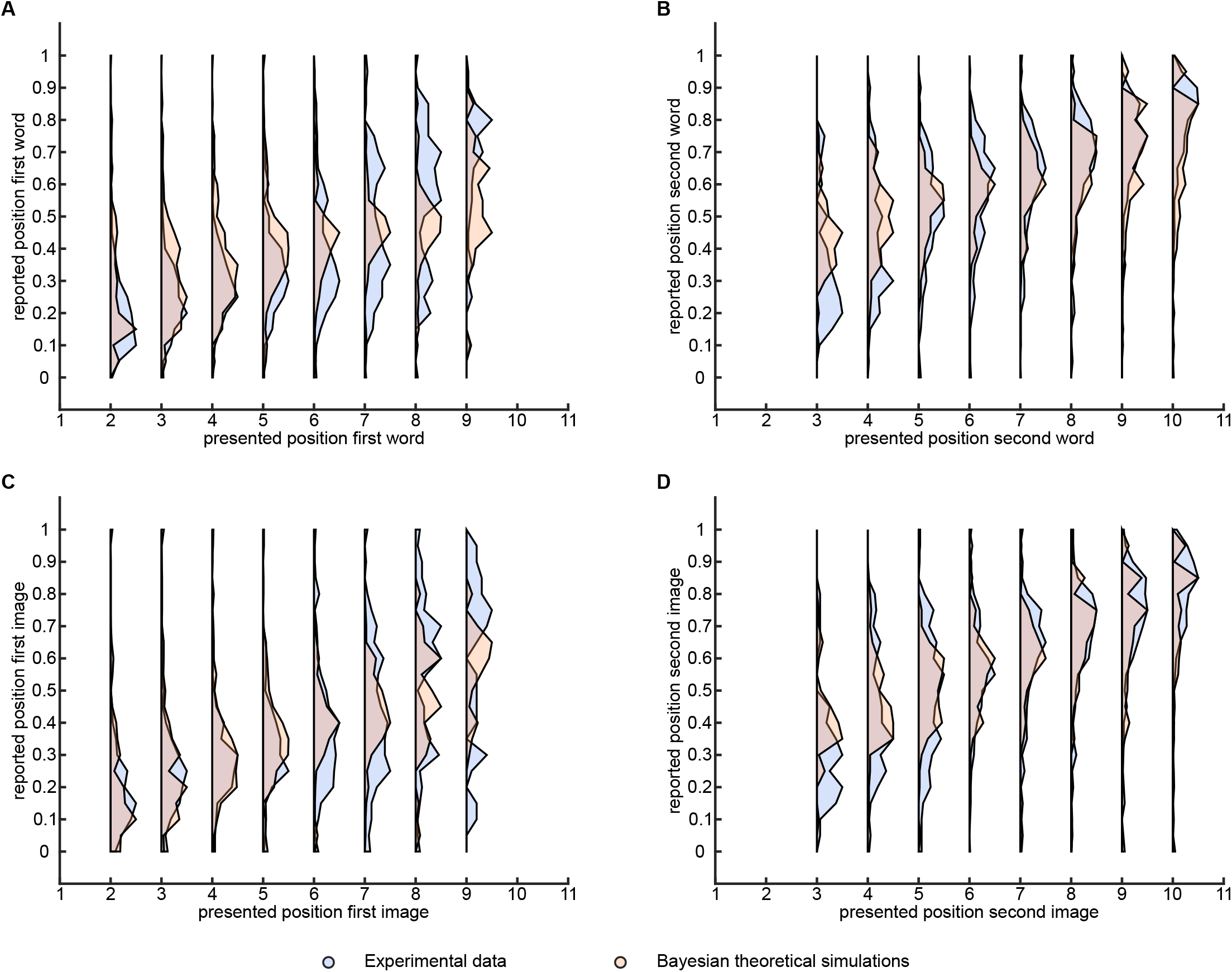
Comparison between Bayesian theory and 2nd experiment. **(A)**: For each presentation time of first intermediate word distribution, normalized by the maximum, of reported times. Blue corresponds to experimental data, while red to theoretical simulations. **(B)**: Same for second intermediate word. **(C)**: Same for first intermediate image. **(D)**: Same for second intermediate image.

**Figure S3:**
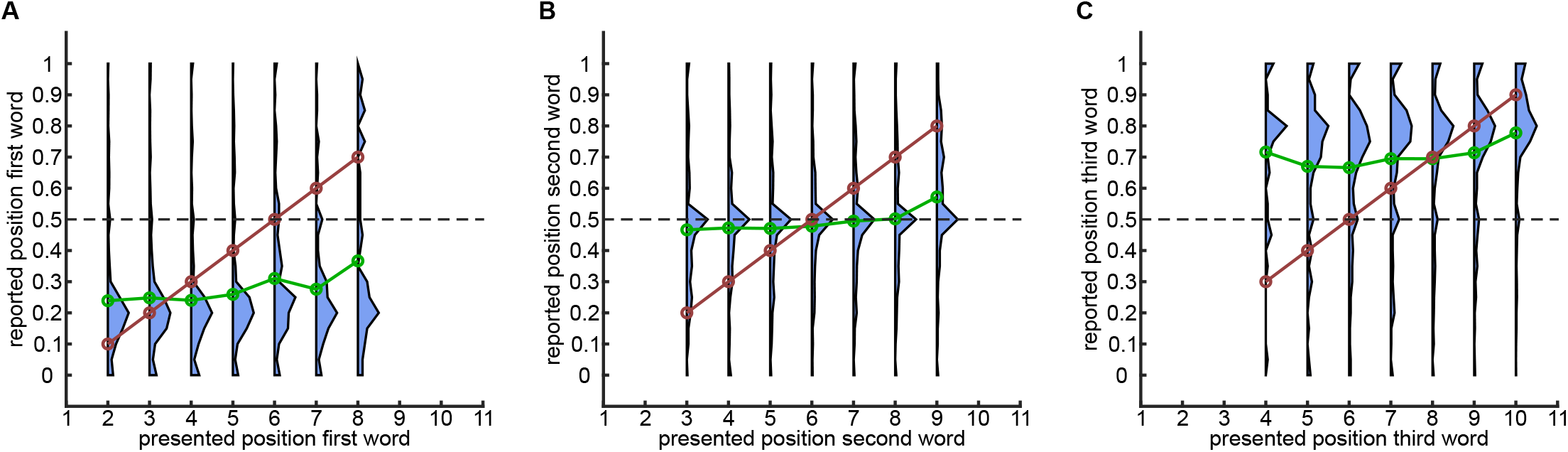
Experiment 3: distribution of reported times. **(A)**: For each presentation time of first intermediate word distribution of reported times. Green line corresponds to average of the distributions, dashed red line corresponds to perfect report. **(B)**: Same for second intermediate word. **(C)**: Same for third intermediate word.

**Figure S4:**
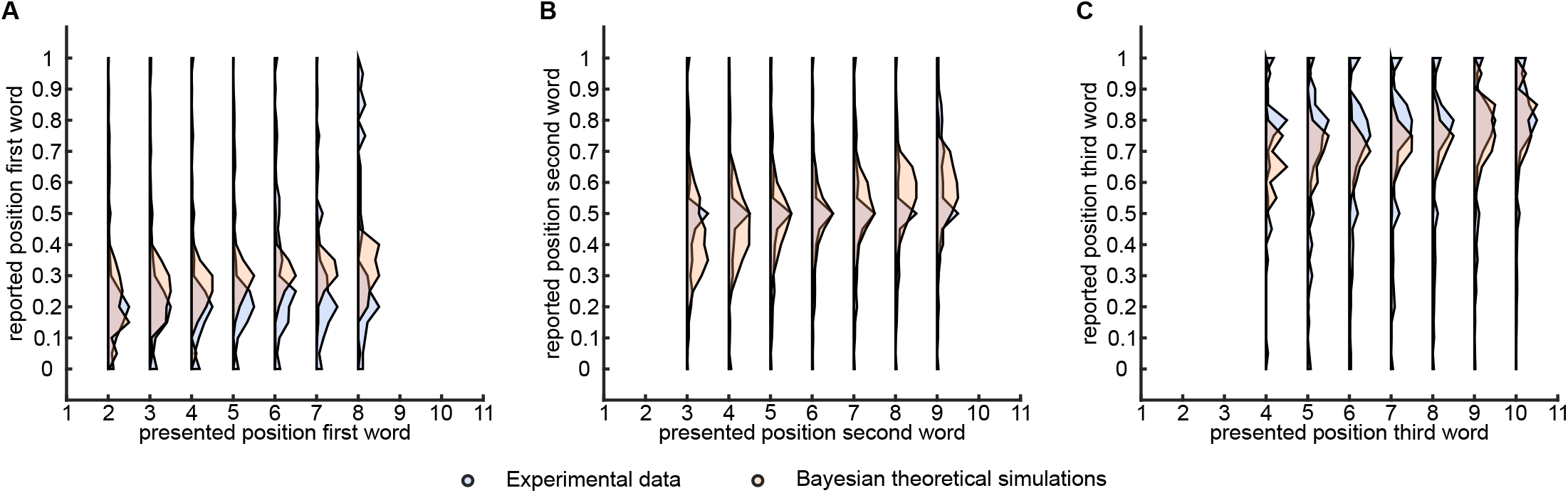
Comparison between Bayesian theory and 3rd experiment. **(A)**: For each presentation time of first intermediate word distribution, normalized by the maximum, of reported times. Blue corresponds to experimental data, while red to theoretical simulations. **(B)**: Same for second intermediate word. **(C)**: Same for third intermediate word.

**Figure S5:**
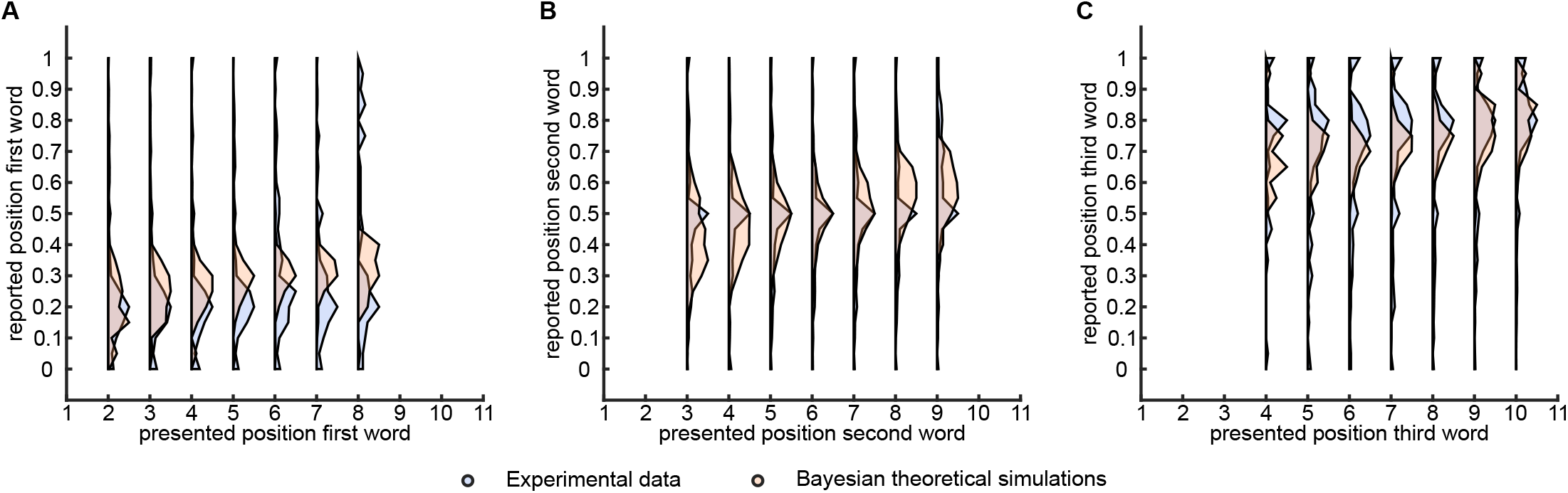
Comparison between Quantized Bayesian theory and 3rd experiment. **(A)**: For each presentation time of first intermediate word distribution of reported times. Blue corresponds to experimental data, while green to theoretical simulations. **(B)**: Same for second intermediate word. **(C)**: Same for third intermediate word.

**Figure S6:**
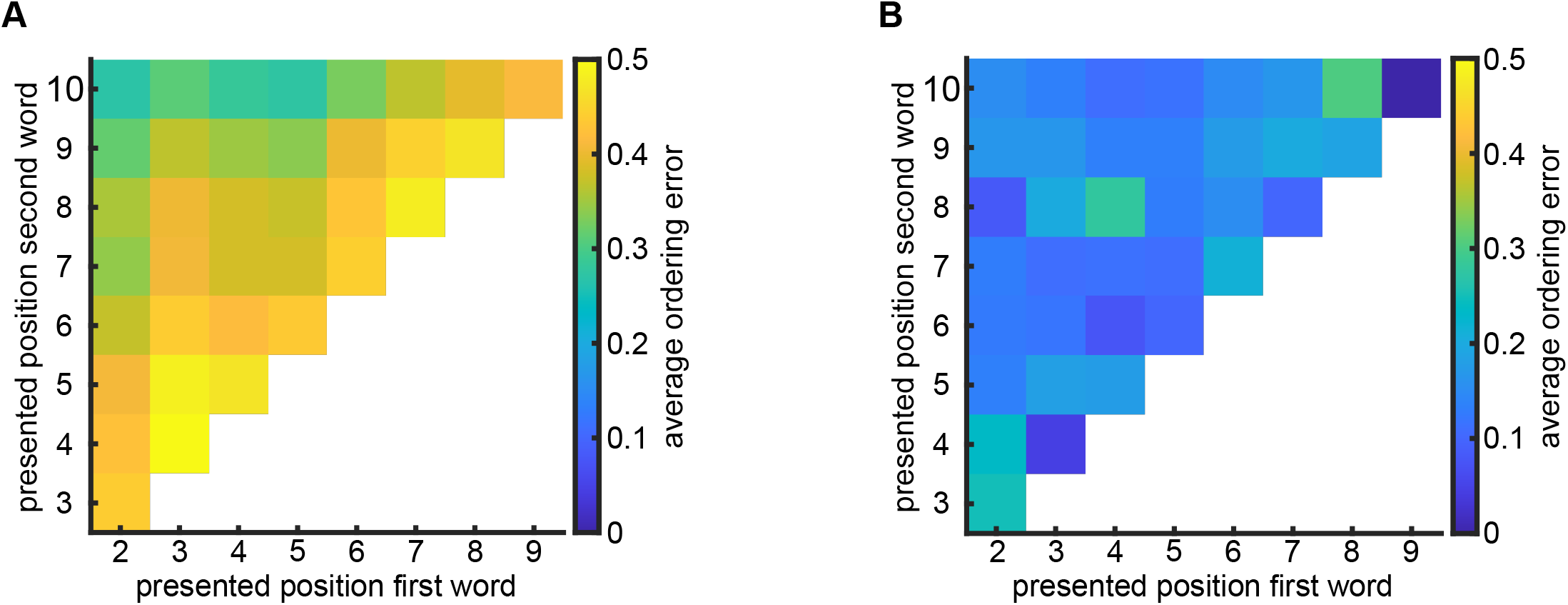
Accuracy of relative time ordering in Experiment 2 with mathematical questions. **(A)**: Naïve prediction of average ordering error from independent distributions obtained in the Experiment 1 with words. **(B)**: Experimental average ordering error with two presented words.

**Figure S7:**
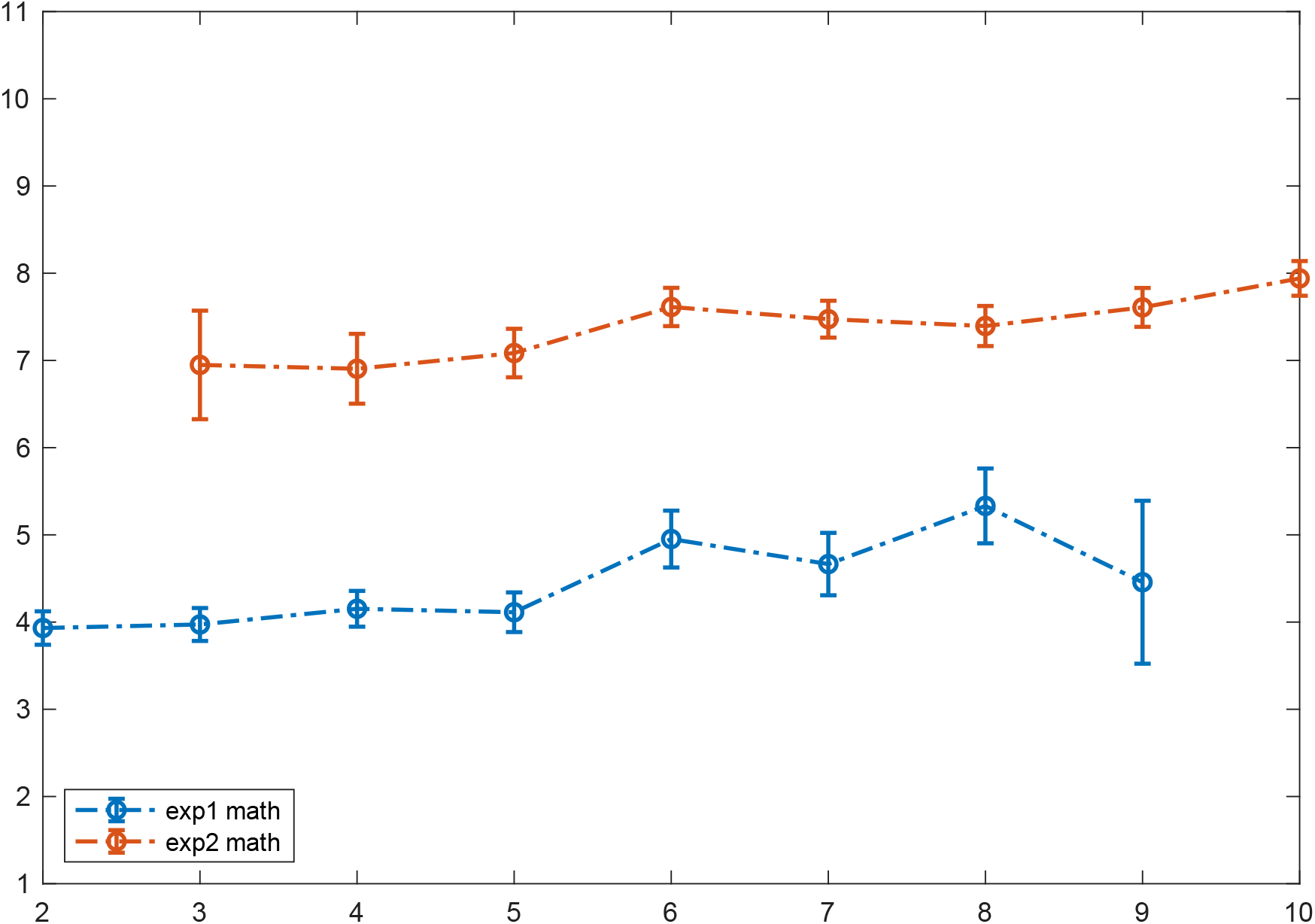
Experiment 4: average time reports. Average report times for first and second word, separately

